# Bisected graph matching improves automated pairing of bilaterally homologous neurons from connectomes

**DOI:** 10.1101/2022.05.19.492713

**Authors:** Benjamin D. Pedigo, Michael Winding, Carey E. Priebe, Joshua T. Vogelstein

## Abstract

Graph matching algorithms attempt to find the best correspondence between the nodes of two networks. These techniques have been used to match individual neurons in nanoscale connectomes – in particular, to find pairings of neurons across hemispheres. However, since graph matching techniques deal with two isolated networks, they have only utilized the ipsilateral (same hemisphere) subgraphs when performing the matching. Here, we present a modification to a state-of-the-art graph matching algorithm which allows it to solve what we call the bisected graph matching problem. This modification allows us to leverage the connections between the brain hemispheres when predicting neuron pairs. Via simulations and experiments on real connectome datasets, we show that this approach improves matching accuracy when sufficient edge correlation is present between the contralateral (between hemisphere) subgraphs. We also show how matching accuracy can be further improved by combining our approach with previously proposed extensions to graph matching, which utilize edge types and previously known neuron pairings. We expect that our proposed method will improve future endeavors to accurately match neurons across hemispheres in connectomes, and be useful in other applications where the bisected graph matching problem arises.

## 1 Introduction

Graph matching is a widely used optimization technique whereby one can find a matching between the nodes in one network and those in another. Solving the graph matching problem yields a matching (i.e. the correspondence between the nodes of the two networks) which minimizes edge disagreements between the two networks. The graph matching problem has found uses in fields as disparate as computer vision [1], biometrics [1], social networks [2], and natural language processing [3], to name just a few.

Most important for this work is the use of graph matching techniques to find bilaterally homologous neuron pairs across the two sides of a nervous system [4, 5]. Connectomes–maps of neural connectivity–can naturally be represented by networks, wherein a node represents a neuron and an edge represents synapses from one neuron to another [6, 7]. Previous works used graph matching techniques to predict neuron pairings between brain hemispheres based on the observed connectivity [4, 5]. The graph matching problem by its very formulation is concerned with two separate networks; as such, previous applications of graph matching to find homologous neuron pairings across hemispheres have considered one network to be the set of nodes and edges within the left hemisphere, and the other network to be defined likewise for the right hemisphere (see Figure 1). In other words, they have only considered the **ipsilateral** connections which connect within a brain hemisphere, and ignored the **contralateral** connections which connect one side of the nervous system to the other. Contralateral connections are quite common in connectomes studied thus far: in subset of the larval *Drosophila melanogaster* (vinegar fly) connectome [8–28], an edge picked at random from the network has about a 35% chance of being a contralateral connection. It is natural to wonder, then, whether these connections can be used to improve automated neuron pairing.

**Figure 1:**
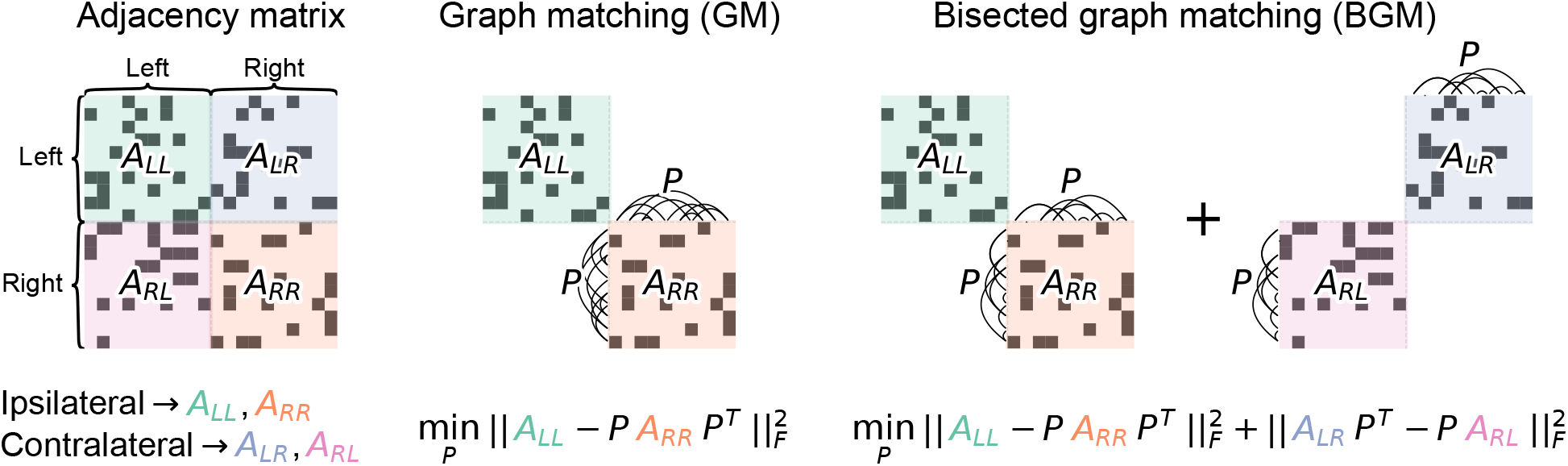
Schematic describing graph matching (GM) and bisected graph matching (BGM). Both aim to find a matching (which can be represented by a permutation, *P*) of the nodes of one hemisphere with respect to the other. GM does so by minimizing the norm of the edge differences between ipsilateral (*A*_*LL*_ and *A*_*RR*_) subgraphs under some matching. BGM aims to jointly minimize the norm of edge differences between ipsilateral *and* contralateral (*A*_*LR*_ and *A*_*RL*_) subgraphs under the same matching applied to both. Note that for the contralateral subgraphs, a permutation of the right hemisphere nodes amounts to permuting the columns (for *A*_*LR*_) *or* the rows (for *A*_*RL*_), but not both.

Here, we show that rather than ignoring the contralateral connections for the purposes of predicting neuron pairs, they can be explicitly included in the optimization by generalizing graph matching to a single network which has been split into two parts. We demonstrate via simulation that when sufficient edge correlations exist between the contralateral subgraphs, our proposed method provides an improvement in matching accuracy. We then show that this methodology indeed improves matching accuracy in our motivating example of a bilateral nervous system by comparing our algorithm to traditional graph matching on five connectome datasets. Further, we describe how our method can be combined with previously proposed generalizations of graph matching to further improve performance.

## 2 From graph matching to bisected graph matching

First, consider the graph matching problem from the perspective of attempting to predict neuron pairs between brain hemispheres (thus adopting the terminology of left/right, etc.), though the techniques described here could be applied more generally. For now, consider the case where both hemispheres have exactly *n* nodes, though methods for graph matching with an unequal number of nodes have been described [29]. Let *A*_*LL*_ be the *n* × *n* adjacency matrix for the subgraph of connections from left hemisphere to left hemisphere neurons, and let *A*_*RR*_ be defined likewise for the right hemisphere. The graph matching problem can then be written as

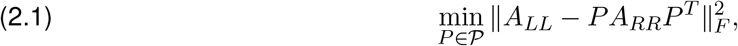

where the set of permutation matrices on *n* nodes is denoted by 𝒫. This objective function measures the number of edge disagreements for an unweighted network, or the norm of the weight disagreements for a weighted network. By trying to minimize this quantity over the set of permutations, one can search for a matching between the networks under which the observed edge structure appears similar.

We are interested in some similar measure which also includes the fact that we want the contralateral connections, under some matching, to appear similar. To formalize this, let *A*_*LR*_ be the adjacency matrix for the subgraph of connections from left hemisphere to right hemisphere neurons, and let *A*_*RL*_ be defined likewise for the connections from the right to the left. We add a term to the graph matching objective function which measures the disagreement between the contralateral subgraphs under some permutation of the nodes of the right hemisphere:

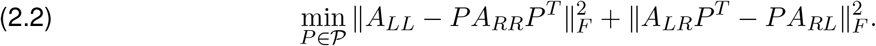

We call the problem in Equation 2.2 the **bisected graph matching problem** (illustrated in Figure 1). With this formulation, the graph matching problem can be seen as a special case of the bisected graph matching problem, since the objective function in Equation 2.2 reduces to that of Equation 2.1 in the special case where *A*_*LR*_ and *A*_*RL*_ are both the zero matrix. Note that this problem is also distinct from the multiplex graph matching problem described in Pantazis et al. [30], as the contralateral subgraphs require only a permutation of their rows *or* their columns (not both) to maintain the correct structure of the adjacency matrix.

Given this notion of what it means to find a good matching between the hemispheres, our goal was to develop an algorithm which could efficiently solve this problem (Equation 2.2). Unfortunately, graph matching problems in general are known to be NP-hard [31], and as such efficient algorithms for solving these problems are approximations. One popular approximation-based algorithm is the Fast Approximate Quadratic (FAQ) algorithm of Vogelstein et al. [32]. This algorithm first relaxes the (discrete) graph matching problem to the relaxed graph matching problem, allowing the tools of continuous optimization to be used (see Section 7.1 for discussion of this approximation). FAQ then uses the Frank-Wolfe method [33] to attempt to minimize Equation 2.1.

The Frank-Wolfe method finds a search direction by minimizing a first-order Taylor series of the objective function, requiring that we compute the objective function’s gradient with respect to its argument, *P*. The gradient of Equation 2.2 with respect to *P* (see Section 11.4 for a derivation) is

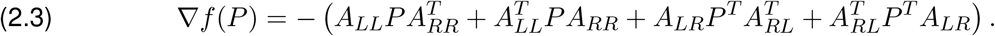

By substituting this new gradient calculation into the FAQ algorithm and keeping the rest of the algorithm the same, FAQ can be adapted to solve the bisected graph matching problem. We provide a full description of the modified FAQ algorithm in Section 7.1. For the remainder of the paper, graph matching (GM) refers to the use of FAQ, while bisected graph matching (BGM) refers to the use of FAQ as modified above.

## 3 Simulation

Here, we demonstrate that this approach improves matching accuracy in simulated datasets when there is sufficient correlation in the contralateral subgraphs. To understand how this correlation affects the usefulness of bisected graph matching, we created simulated data using the correlated Erdős-Rényi model (*CorrER*). The correlated Erdős-Rényi model is a special case of the correlated stochastic block model introduced in Lyzinski et al. [34]. Briefly, a pair of networks is distributed *CorrER*(*n, p, ρ*) if both networks marginally are distributed as Erdős-Rényi models [35, 36] with *n* nodes and global connection probability *p*, but the edges of the two networks have Pearson correlation *ρ*. Note that this correlation of edges also requires specifying an alignment of one network to the other, which we can use as ground truth for evaluating our algorhtm. Here, we use the version of this model for a directed network to more closely resemble nanoscale connectome data, which has directed edges. We used the correlated Erdős-Rényi model (as implemented in graspologic [37]) to construct a simulation of a “bilateral” network as follows:

1. The ipsilateral subgraphs were sampled from a correlated Erdős-Rényi model (*CorrER*):

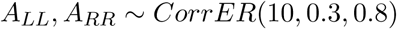
2. Independently from the ipsilateral networks, the contralateral subgraphs were sampled from a correlated Erdős-Rényi model:

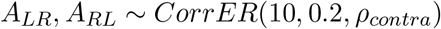
3. The full network was defined as

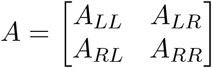
4. To simulate an unknown correspondence between the nodes of the left and right hemispheres, we applied a random permutation (*P*_*rand*_) to the nodes of the “right hemisphere” in each sampled network:

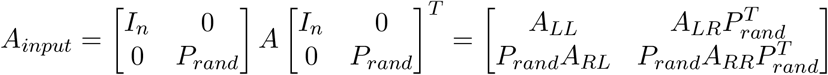

*A*_*input*_ is the network which was input to the matching algorithms, obscuring the true matching from the true alignment in order to evaluate their performance.

We varied the value of *ρ*_*contra*_ from zero (*A*_*LR*_ and *A*_*RL*_ have no correlation, and thus are not helpful for matching) to one (*A*_*LR*_ and *A*_*RL*_ are isomorphic, providing extremely helpful information for matching). For each value of *ρ*_*contra*_, we simulated 1,000 networks. For each network, we ran the graph matching (GM) and bisected graph matching (BGM) algorithms to attempt to uncover the correct permutation which would realign the left and right hemispheres. For both algorithms, we used default parameters, and one initialization for each algorithm. For each run of each algorithm, we examined the matching accuracy, which is the proportion of nodes correctly matched.

Figure 2 shows the matching accuracy for both algorithms as a function of *ρ*_*contra*_. For low values of *ρ*_*contra*_, using bisected graph matching actually degrades performance. When *ρ*_*contra*_ = 0, the match accuracy drops by ∼29%. This is unsurprising, as in this case the contralateral subgraphs are effectively noise with respect to the correct matching between the left and right. For small values of *ρ*_*contra*_, bisected graph matching often found permutations of the contralateral subgraphs which had *fewer* edge disagreements than the alignment used to generate the correlated networks (Section 11.1), explaining why including these connections pulls the solution away from the true matching. However, as *ρ*_*contra*_ increases, bisected graph matching eventually outperforms graph matching. For this simulation, when *ρ*_*contra*_ is greater than 0.4, the accuracy for bisected graph matching is higher (by more than ∼20% when *ρ*_*contra*_ ≥ 0.9). We also found that this phenomenon was consistent across a range of network sizes (Section 11.2).

**Figure 2:**
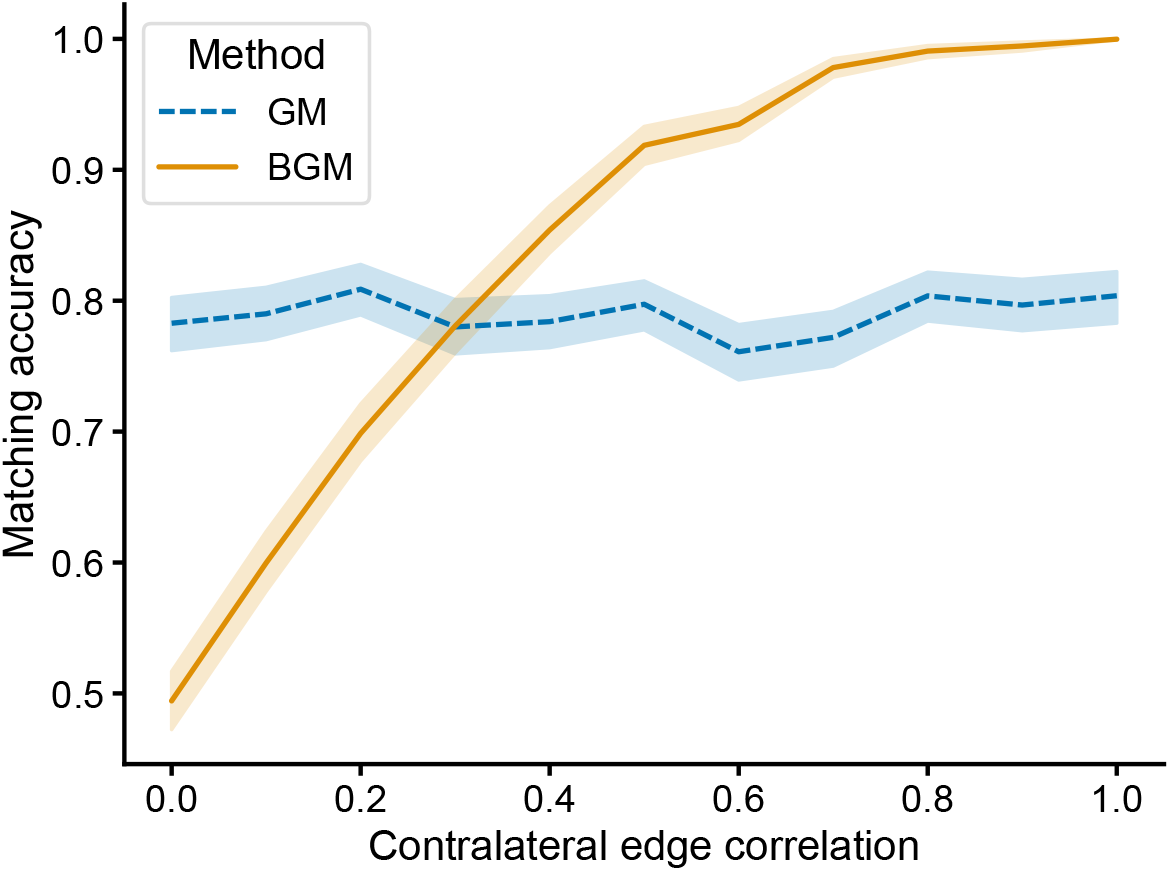
Performance of graph matching (GM) using the FAQ algorithm [32] and bisected graph matching (BGM) (this work) on a simulated dataset constructed such that the ipsilateral and contralateral connections both come from correlated Erdős-Rényi models (Section 3). Each network had 10 nodes per side, the ipsilateral connection density was 0.3, the ipsilateral edge correlation was 0.8, and the contralateral connection density was 0.2. We varied the contralateral edge correlation from 0 to 1, and for each value, we simulated 1000 networks and ran both algorithms on the same data with the same initialization. Lines show the mean matching accuracy, shaded regions show 95% confidence intervals. As the correlation in the contralateral connections increases, including them in the optimization becomes more helpful.

Whether bisected graph matching improves accuracy is determined by many factors, including the correlation in contralateral edge structure studied here in this simple simulation. We next sought to see whether this bisected graph matching would be helpful in our motivating example of matching neurons between two sides of a nervous system.

## 4 Connectomes

We examined the performance of both graph matching algorithms on a set of real connectome datasets. To ensure we could evaluate the performance of both algorithms, we restricted our analysis to connectomes for which pairings of individual neurons between sides of the nervous system were already known. We studied the (chemical) connectomes of both a hermaphrodite and a male *Caenorhabditis elegans* worm [39], the pharynges of two *Pristionchus pacificus* worms [38], and a subset of a larval *Drosophila melanogaster* [8–28]. For all these datasets, neuron pairings across sides of the nervous system are not complete - indeed, some neurons appear only on one side of the organism or exactly in the center [39]. Thus, we restricted our analysis to the subset of neurons which were present as a bilaterally homologous pair and for which this pairing was known. We then ensured that the remaining set of nodes was fully (weakly) connected for each dataset, removing nodes not part of the largest connected component. Table 1 shows the number of nodes and edges remaining for each of the connectome datasets considered here. We treat each network as weighted (by synapse count) and directed (since the direction of chemical synapses is known).

**Table 1:** Summary of the connectome datasets studied in Section 4, showing the number of nodes, number of edges, percentage of contralateral synapses, correlation of ipsilateral subgraphs, and correlation of contralateral subgraphs for each connectome. Correlations are Pearson’s correlation coefficient. All metrics are with respect to the datasets after processing to select fully connected networks composed of neurons who have bilateral pairs, as described in Section 4. The number of nodes per hemisphere and (and the number of known pairs) is always half the total number of nodes, since only paired neurons are considered.

| Dataset | # nodes | # edges | % contra.<br>synapses | Ipsi.<br>correlation | Contra.<br>correlation |
| --- | --- | --- | --- | --- | --- |
| *P. pacificus* pharynx 1 [38] | 18 | 35 | 30 | 0.79 | 0.83 |
| *P. pacificus* pharynx 2 [38] | 22 | 33 | 32 | 0.84 | 0.88 |
| *C. elegans* hermaphrodite [39] | 286 | 2,838 | 41 | 0.87 | 0.86 |
| *C. elegans* male [39] | 360 | 2,482 | 37 | 0.81 | 0.70 |
| *D. melanogaster* larva subset [8–28] | 1,240 | 32,564 | 31 | 0.88 | 0.83 |

For each connectome, we predicted each neuron’s pair on the other side of the nervous system by applying either the graph matching or bisected graph matching algorithms. We ran 50 initializations, each from the barycenter, since neither algorithm is deterministic (see Section 7.1 for more explanation on initialization and randomness in the algorithm). For each initialization, we ran both algorithms with default parameters [37], and measured the matching accuracy with respect to the known pairing of neurons.

We observed that for all the connectomes studied here, bisected graph matching improves matching performance (Figure 3), sometimes dramatically so. For all five connectomes, the match ratio for the bisected graph matching algorithm was significantly higher (*p <* 0.0005 for each dataset, two-sided Mann-Whitney U tests). Matching accuracy increased by ∼44% and ∼20% for the two *P. pacificus* samples, ∼29% for the hermaphrodite *C. elegans*, ∼15% for the male *C. elegans*, and ∼22% for the *Drosophila* larva subset, respectively. We also found that this trend holds when we relaxed the requirement that all nodes in the connectome have a homologous partner, finding that bisected graph matching always provided increased matching accuracy even when some neuron pairs in these connectomes were artificially unmatched (Section 11.3).

**Figure 3:**
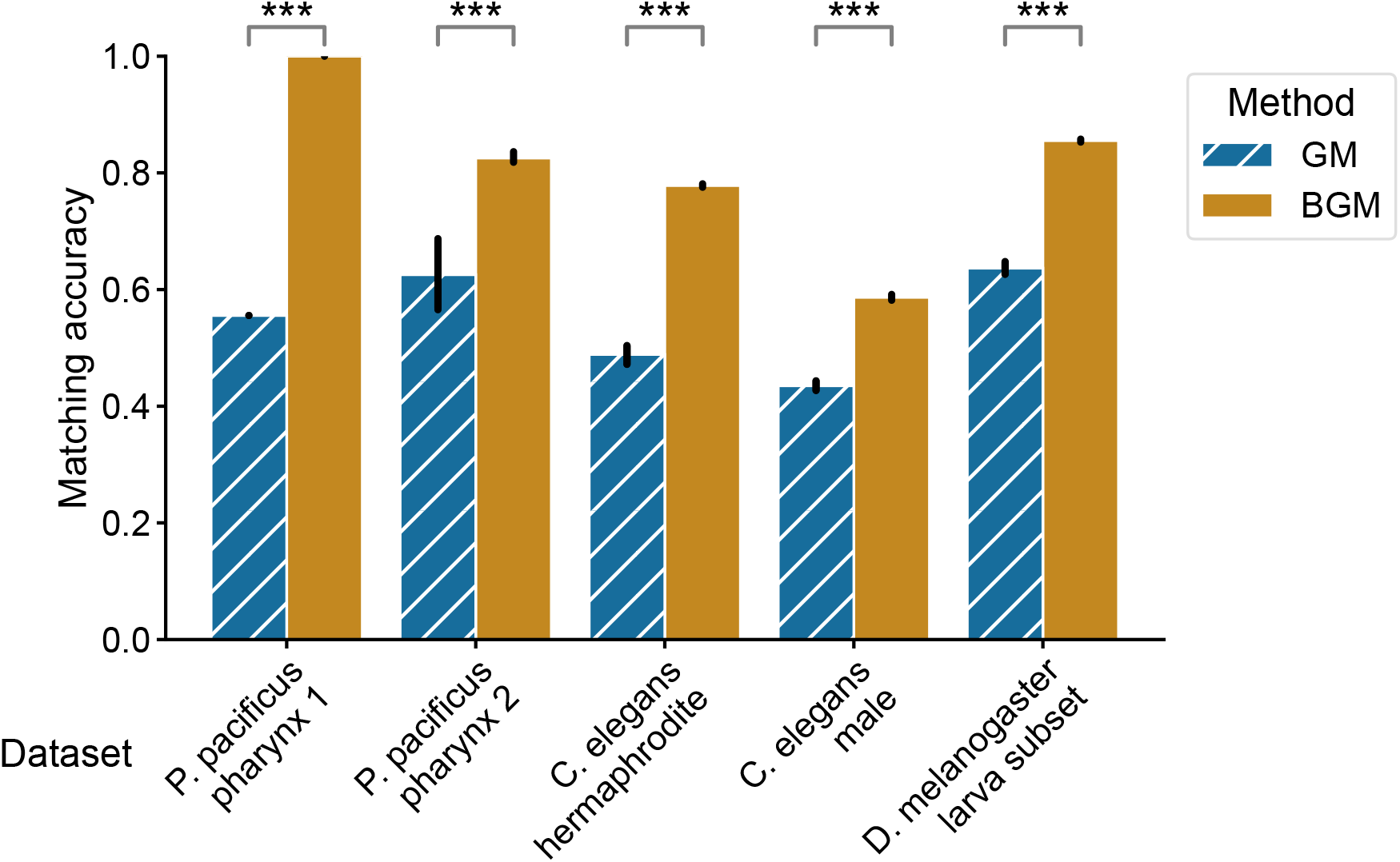
Performance of graph matching (GM) and bisected graph matching (BGM) on five bilateral connectome datasets. For each comparison, we performed 50 initializations, and from each initialization we ran both the graph matching and the bisected graph matching algorithms. The mean match accuracy is shown for both methods on each dataset. Error bars show 95% confidence intervals for the mean. For every dataset, bisected graph matching provided a performance improvement over graph matching which ignores contralateral connections. *** indicates a significant difference where *p <* 0.0005 (two-sided Mann-Whitney U test).

These results demonstrate the practical utility of our proposed algorithm for improving bilaterally homologous pair prediction in connectomes. We next sought to show how our proposed method can be combined with previously described extensions of graph matching to further improve performance.

## 5 Extensions

### 5.1 Matching multiplex networks

While connectomes are often described as networks, many of these datasets actually lend themselves to multiplex network representations. For the purposes of this paper, we consider multiplex networks to have one set of nodes, but potentially multiple types of edges between these nodes (see for Kivelä et al. [40] for a review of multilayer networks more generally). For instance, in *C. elegans*, both chemical (synaptic) and electrical (gap junction) connections have been mapped [39]. If we consider these connections to each be of their own “type,” then we can construct an adjacency matrix for each– these become the “layers” of our multiplex network. As further examples of edge types in connectomics, *Drosophila* connectomes are beginning to have neurotransmitter information associated with each synapse [41], as well as a differentiation between axo-axonic, axo-dendritic, dendro-axonic, and dendro-dendritic connections [42, 43].

To match neurons based on connectivity using this multiplex network information, one can generalize the graph matching problem to a multiplex graph matching problem. Pantazis et al. [30] proposed a generalization of the FAQ algorithm to solve this problem. This multiplex graph matching scheme can easily be combined with the bisected graph matching proposed in this work, again by simply modifying the graph matching objective function and its gradient to account for these multiple connection types (see Section 7.2 for more details).

We applied graph matching and bisected graph matching to the connectomes of both *C. elegans* sexes, and varied whether the networks used were either chemical, electrical, or both (multiplex network). Figure 4 displays matching accuracy for both algorithms using each combination of edge types. We observed a clear advantage to using multilayer graph matching on these datasets: in both connectomes and for both GM and BGM, matching with the multilayer network outperformed matching for either chemical or electrical connections alone. We also found that BGM outperforms GM for any combination of network layers for both connectomes. These results highlight the advantages of combining BGM with a previously described extension of graph matching [30] when multiple edge types are available, as the highest accuracy on both datasets came from using both contralateral connections and multiple edge types.

**Figure 4:**
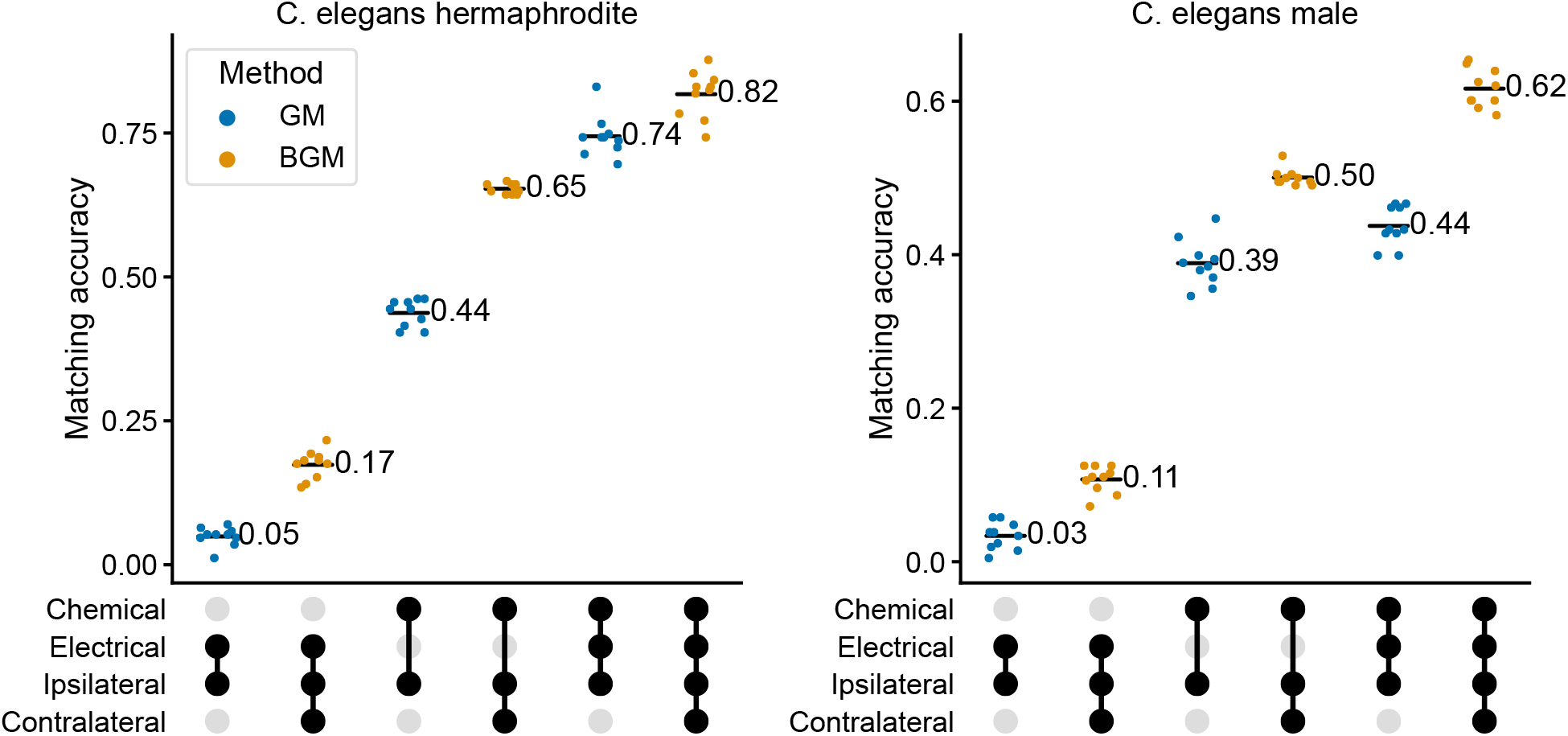
Matching accuracy on the hermaphrodite (left) and male (right) *C. elegans* connectomes. Accuracy is shown when using various combinations of network layers (chemical, electrical, or both) and subgraphs (ipsilateral (GM) or ipsilateral and contralateral (BGM)). Labels denote the mean matching accuracy over 10 initializations. Filled circles below the x-axis indicate the layers/subgraphs used for matching in a given column. For each combination of layers, BGM always showed an increase in mean matching accuracy over GM (p-values *<* 0.005 for all of these comparisons, two-sided Mann-Whitney U test). On both datasets, the best results came from using BGM (this work) in concert with the multiplex graph matching proposed in Pantazis et al. [30].

### 5.2 Networks of differing sizes and with seeds

Next, we studied how BGM would work with two further extensions to graph matching based on the work of Fishkind et al. [29]. Often, the two hemispheres being matched may not have exactly the same number of neurons, but one still wishes to find a matching between them. Further, partial matching information is also common – for instance, one could have complete matching information about a subset of nodes in some brain region, and would like to use this partial matching to improve matching of the rest of the brain. Fishkind et al. [29] studied exactly this setting, proposing “padding” schemes to deal with networks which have different sizes, as well as a method for incorporating a partial matching or “seed” nodes into the FAQ algorithm.

We applied the corresponding generalizations of these ideas to the bisected graph matching case (see Section 7.3 for details), allowing us to apply our proposed algorithm to a dataset where a partial matching was known ahead of time and the number of neurons in the two hemispheres was not the same. To demonstrate these capabilities, we applied this method to the *Drosophila* larva partial connectome [8–28]. In Section 4, we restricted our analysis to the set of nodes for which published pairings existed, such that we could evaluate matching accuracy. Here, we relaxed this restriction, and used these published pairs as seed nodes. We also note that the full collection of published neurons has 942 neurons on the left hemisphere and 938 neurons on the right hemisphere. The padded graph matching of Fishkind et al. [29] allowed us to perform a matching on these two networks of differing sizes (resulting in some neurons on the larger hemisphere not being matched in each run of the algorithm).

To examine the effect of seeds, we performed a cross-validation-like experiment, wherein some seeds (20%) were reserved for evaluation so that we could compute matching accuracy. We used some number of the remaining pairings as seeds for either graph matching or bisected graph matching. Figure 5 shows matching accuracy on these held-out known pairings as a function of the number of seeds used. We found that for any number of seeds, bisected graph matching always had a higher mean matching accuracy than graph matching. Conversely, BGM can be viewed as allowing the user to reach the same accuracy level for a smaller number of previously known seeds, which can be effortful to obtain for a new dataset.

**Figure 5:**
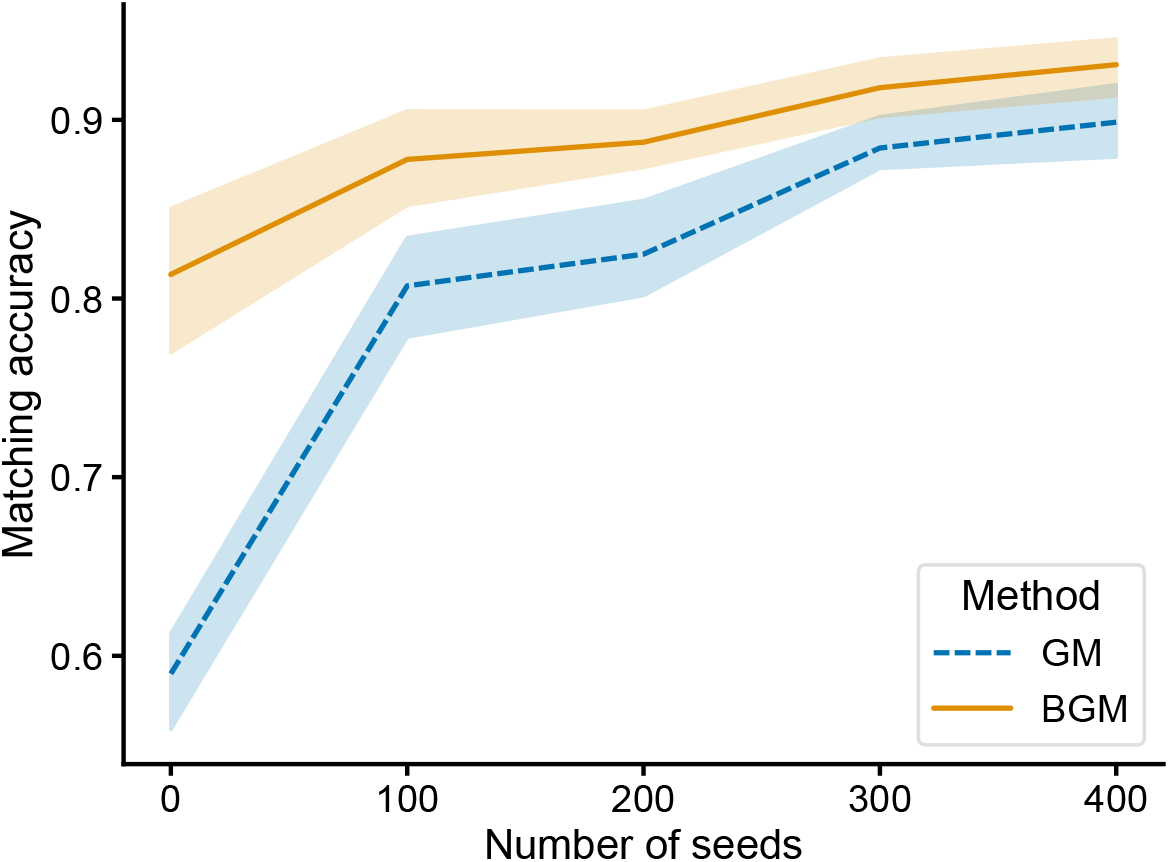
Matching accuracy using seeded matching techniques on the *Drosophila* larva connectome subset [8–28]. Here, both GM and BGM leveraged the previously published paired neurons as “seeds” which can be used to improve the matching of the rest of the network [29]. Average matching accuracy is shown across 5-folds of cross-validation: 20% of seeds were used for evaluating accuracy, and some number of the remaining seeds (x-axis) were input to GM and BGM. Regardless of the number of seeds, BGM always provided an accuracy improvement.

Given the superiority of BGM over GM across a range of experiments, we finally sought to examine the matches for neurons where we did not know of a previously presented pairing. We re-ran 100 initializations of bisected graph matching on the *Drosophila* larva subset, using all known pairings as seeds. Figure 6 shows the morphology of 6 example predicted neuron pairs which were always matched together across all 100 initializations. We found that the morphology of these frequently matched neurons was generally similar, suggesting that they may represent true bilaterally homologous pairings. Further investigation will be required to confirm or reject these candidate matches, but our results demonstrate how bisected graph matching can be used to easily provide well-informed guesses for these pairings. We also provide examples of neurons which were very *infrequently* paired across each initialization (implying a lower confidence in these matches [29]), suggesting that these pairs are less likely to be true homologs (Supplemental Figure 4). We include all matching results for these previously unpaired neurons in the supplemental information.

**Figure 6:**
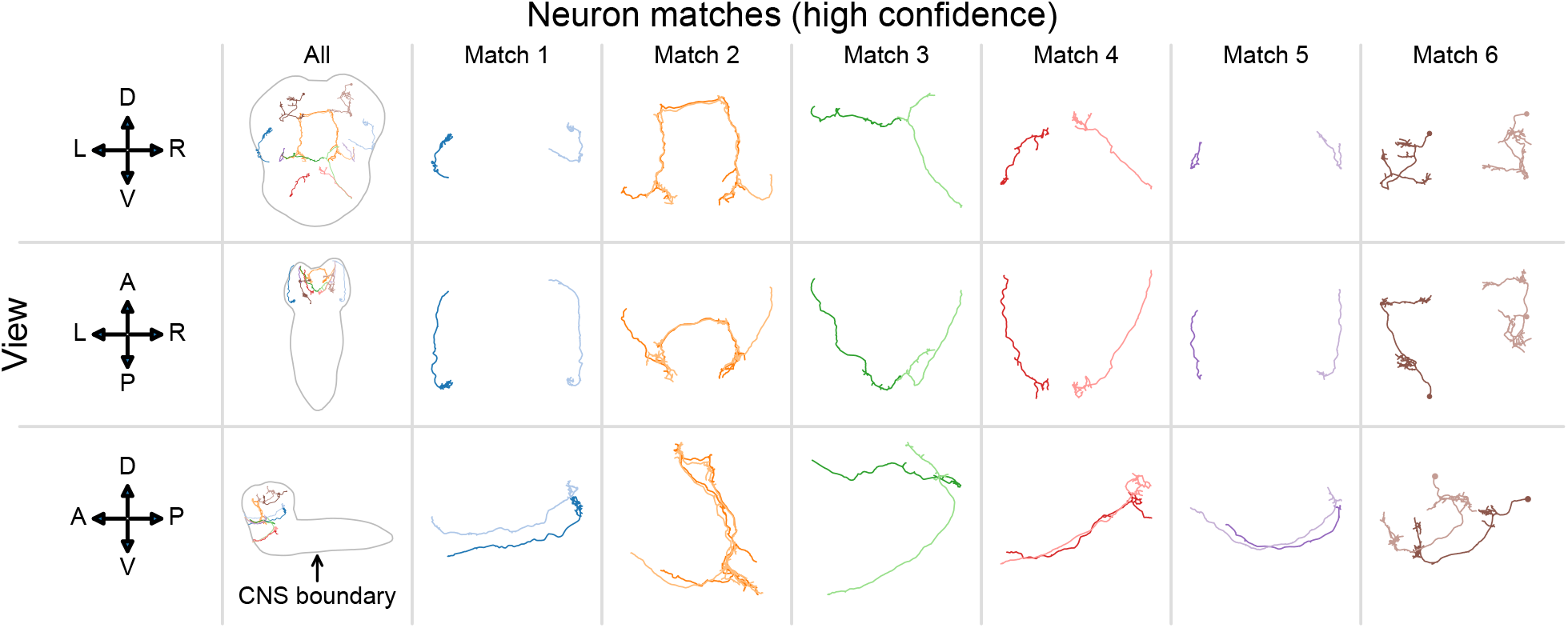
Morphological comparison of matched neurons in the *Drosophila* larva connectome subset [8–28] using all available seeds and the BGM algorithm. Each column shows an example neuron match which was always selected by BGM across 100 initializations, indicating high confidence in that match [29]. Each row shows a different view of a matched pair of neurons (anatomical axes to the left show: D-dorsal, V-ventral, L-left, R-right, A-anterior, P-posterior). The morphology of these matched neurons appears similar, suggesting that these are plausible candidates for previously undescribed bilaterally homologous neurons.

## 6 Discussion

### 6.1 Summary

We proposed a simple generalization of the graph matching problem, which incorporates any connections *between* the two sets of nodes being matched. We then showed how this problem could be solved by using a new objective function in the framework of a state-of-the-art graph matching algorithm, FAQ [32]. In simulations, we saw that as the strength of the correlation between the contralateral subgraphs increases, these connections become more useful to include in the matching process. By running both graph matching and bisected graph matching on five connectome datasets, we provided compelling evidence that for practical purposes in neuroscience, including these contralateral connections in the optimization is beneficial. We further showed how our algorithm can be applied to settings involving multiplex networks, networks of differing sizes, and how the algorithm can leverage a partial, known pairing of neurons to improve matching performance for the remaining neurons. We have provided a documented, open-source implementation of our algorithm (Python 3) to enable its easy application to future connectome datasets (see Section 8).

### 6.2 Limitations

As we showed in simulation in Section 3, bisected graph matching is only likely to improve matching accuracy in connectomes when there is sufficient correlation between the contralateral subgraphs. For a new organism (or possibly even just a new sample), this will not be known in practice. Domain knowledge as to the nature of the contralateral connections in an organism’s brain may be important when choosing whether to include them in the matching as described in this work, though we note that all five connectomes studied in this work had a high (>0.7) contralateral correlation (Table 1). In practice, it may be best to evaluate different matching algorithms (and hyperparameters) on a subset of the connectome prior to matching a complete dataset.

Further, other approaches to matching neurons in neuroscience do not use connectivity at all: Costa et al. [44] introduced an algorithm for matching neurons on the basis of morphology, which has been widely used on connectomic reconstructions. In practice, it is likely that the best neuron pairings will be achieved by a joint optimization which considers morphology, multiple edge types, seeds, and the contralateral connections as proposed in this work. We did not explore this possibility here, but it remains an intriguing future pursuit.

### 6.3 Outlook

As more connectomes are mapped both from new organisms or for more individuals of the same species, the tools provided here will accelerate the process of finding correct pairings of neurons between the two sides of a nervous system, while requiring less human labor to annotate pairs by hand. These neuron pairings appear to be a fundamental property of the invertebrate nervous systems studied in connectomics thus far. Finding these neuron pairs is important for understanding the stereotypy in an organism (i.e., how similar is the connectivity of the left and the right) [39, 45–47]. Additionally, these neuron pairs can be useful for statistical approaches which leverage a one-to-one correspondence of nodes across networks [48, 49]. We also note that the bisected graph matching problem (and the analogous version of the quadratic assignment problem, which is equivalent to the graph matching problem up to a sign change [32]) may arise in other settings where one wishes to match nodes in a single graph which can be split into two parts and some level of symmetry exists between them.

## 7 Methods

### 7.1 Graph matching and bisected graph matching

Since graph matching is an NP-hard problem [1, 31], no efficient algorithm exists which will always yield a perfect matching. Part of the difficulty of this problem is that the search space of permutations is large (there are *n*! permutations of *n* nodes) and discrete (there is no way to interpolate between two permutations and still have a permutation). Thus, many algorithms relax this constraint, enabling efficient solutions of the relaxed problem [32, 50–52]. A common approach (used by FAQ [32] and other algorithms [51, 52]) is to relax the (discontinuous) search for a permutation matrix to the convex hull of this set, the set of doubly stochastic matrices, 𝒟 [32]. Note that Lyzinski et al. [50] showed that under a model of correlated random Bernoulli graphs, the best solution to the relaxation used by FAQ almost always yields the correct permutation matrix as the size of the networks grows (this does not say that FAQ will always find this solution, however).

In this relaxed space of doubly stochastic matrices, FAQ requires an initial position to start its search. A common default value (which we use in this work) is the barycenter, which is the centroid of the set of all doubly stochastic matrices, and is simply an *n* × *n* matrix where all elements are 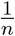. FAQ then proceeds by using the Frank-Wolfe method to iteratively update its search for a doubly stochastic matrix which maps one adjacency matrix to another. The algorithm terminates after either a maximum number of iterations or when the search positions change very little (less than some tolerance parameter) between iterations. After this doubly stochastic solution has been found, FAQ then projects back onto the set of permutation matrices by solving the linear assignment problem.

Below, we detail the BGM-via-Frank-Wolfe algorithm (referred to simply as BGM in the text), which simply adapts this procedure by replacing the objective function and its gradient to solve the bisected graph matching problem as described in Section 2. We refer the interested reader to Vogelstein et al. [31] for further details on the original algorithm, and to Fishkind et al. [29] for many interesting extensions. We also note that implementations of the FAQ algorithm are available in SciPy [53] and graspologic [37].

Two nuances of this algorithm for practical usage are worth commenting on. First, we note that FAQ is not guaranteed to find the correct solution to the graph matching problem (and again, neither is any polynomial-time algorithm [1, 31]). Even if the minimizer to the indefinite relaxed graph matching problem is the correct permutation (as described in Lyzinski et al. [50]), the Frank-Wolfe method may get stuck in a local minima, and not find this best solution. Second, this algorithm is not deterministic—different initializations can lead to different solution paths, which may get stuck in local minima. Even from the same initialization, there may be more than one-step direction (see Algorithm 1 Step 2) at any given position in the solution space, since multiple step directions can be deemed equally suitable. Our implementation simply chooses one of these at random. Thus, even from the same initialization, it is often beneficial to restart the algorithm multiple times, and choose the solution with the best objective function value. For this reason, a number of the experiments in the main text specify the number of initializations used.

#### Algorithm 1 Bisected Graph Matching (BGM) via Frank-Wolfe. To recover the FAQ algorithm [32] (GM), simply set *A*_*LR*_, *A*_*RL*_ to the zero matrix

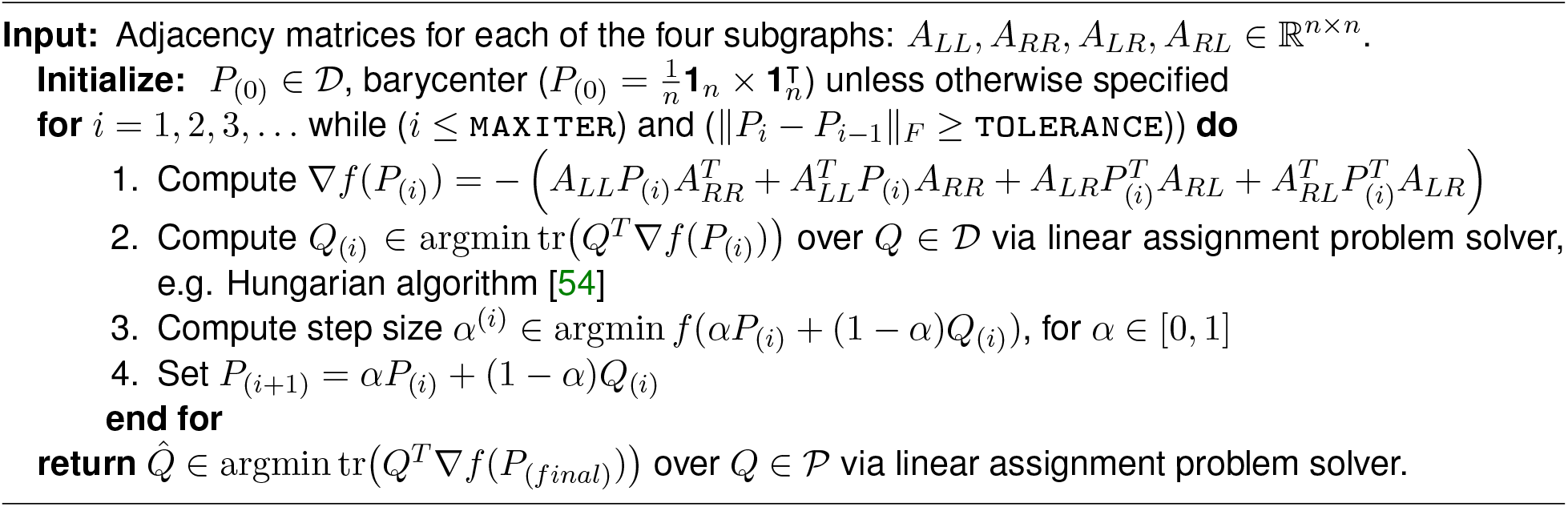

### 7.2 Multilayer graph matching

In this work, we consider a multiplex network to have multiple edge types. If there are *K* different edge types, then a multiplex network (say, *A*_*LL*_) can be represented by *K* different adjacency matrices,

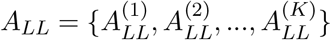

Matching *A* to some other multiplex network (which has the same edge types 1, …, *K*), *A*_*RR*_, thus amounts to matching each of their constituent adjacency matrices. Pantazis et al. [30] formalized this notion by writing the objective function as

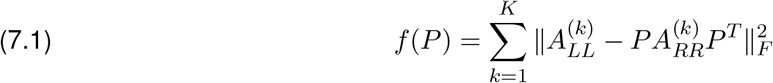

Note that the same permutation matrix, *P*, jointly maps each of these adjacency matrices together. To perform multiplex bisected graph matching, we apply the same generalization to the bisected graph matching objective function (Equation 2.2)

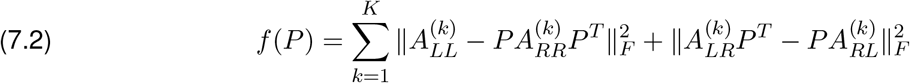

The gradient of this new objective function is simply the sum of the gradients of each term

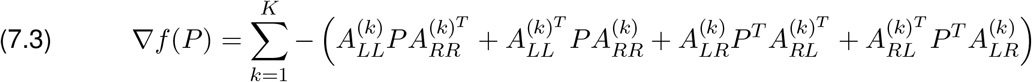

Using Equations 7.2 and 7.3 as the objective and gradient, respectively, in Algorithm 1 yields a method for solving a multiplex bisected graph matching problem.

### 7.3 Seeded graph matching

Fishkind et al. [29] considered modifying FAQ to solve the so-called seeded graph matching problem, wherein a subset of the nodes of the two networks are matched ahead of time. The goal is to leverage these previously known pairings to improve the pairings of the rest of the network. This seeded graph matching problem can be thought of as restricting the search space of all permutations to only those which respect a particular seed set.

We briefly present the methods for seeded graph matching here, and refer the interested reader to Fishkind et al. [29] for more details. Adapting notation to match that of this work, denote the adjacency matrix of seeded-to-seeded connections in the left-to-left subgraph as 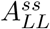, the matrix of seeded-to-nonseeded connections in the left-to-left subgraph as 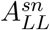, and likewise for the other possible sub-graphs. With this definition, Fishkind et al. [29] showed that the seeded graph matching objective function has the same minimizer as

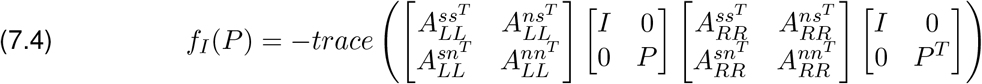

Further, they showed that the gradient of *f*_*I*_(*P*) with respect to *P* is

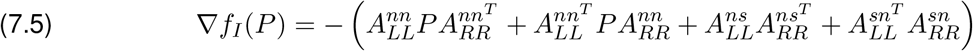

For the term we added to the graph matching objective for the contralateral adjacency matrices, the minimizer is the same as that of

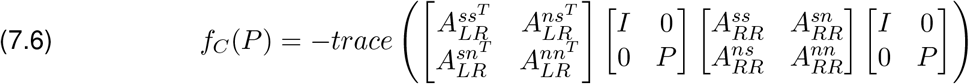

And similarly to the standard graph matching case, the gradient is

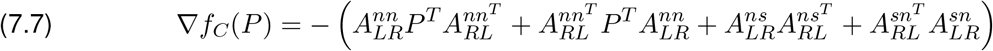

For seeded graph matching with ipsilateral and contralateral connections, the full objective function is *f*_*I*_(*P*) + *f*_*C*_(*P*), and its gradient is ∇*f*_*I*_(*P*) + *f*_*C*_(*P*). We use this new objective function and gradient in the Frank-Wolfe method (Algorithm 1) to yield a seeded bisected graph matching algorithm.

### 7.4 Padded graph matching

Consider the case where *A*_*LL*_ has more nodes than *A*_*RR*_ (without loss of generality, because we could swap the left and right sides to yield an equivalent algorithm). Let *n*_*L*_ be the number of nodes on the left, and *n*_*R*_ be the number of nodes on the right. Fishkind et al. [29] proposed a “padding” scheme, wherein these networks can be made comparable for matching. Their “naive” padding scheme simply replaces *A*_*RR*_ with a new matrix that has added zeros to make it match the size of *A*_*LL*_:

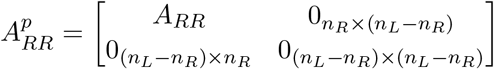

where 0_*m×n*_ is a *m* times *n* matrix of all zeros. 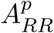 can now be matched to *A*_*LL*_, though some nodes on the left would be matched to row/columns of all zeros, and therefore not have a valid match on the right.

For bisected graph matching, we use the same padding idea adapted to our setting. The padded version of *A*_*LR*_ is

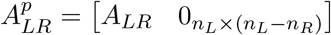

and the padded version of *A*_*RL*_ is

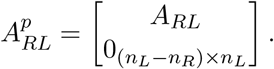

For padded bisected graph matching, these matrices 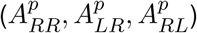 are used in place of the original subgraphs such that the graph matching algorithms described above (which require matrices to be of the same size) can be applied. We did not explore the use of the “adopted” padding scheme, as in our case the two sides of the connectome being matched had approximately the same number of nodes, and this method was not described for weighted networks [29].

## 8 Code and data

Analyses relied on graspologic [37], NumPy [55], SciPy [53], Pandas [56], NetworkX [57], and pymaid [58]. Plotting was performed using matplotlib [59], Seaborn [60], and NAVis[61].

All code for this paper (implemented in Python 3) can be found on GitHub at github.com/neurodata/bgm and viewed as a Jupyter Book [62] at docs.neurodata.io/bgm. There are no primary data in the paper (see references in the main text Table 1). All data is included in the GitHub repository. The source code and data is also archived at doi.org/10.5281/zenodo.6561550.

## 9 Author contributions

- Conceptualization: B.D.P., M.W., C.E.P., J.T.V.
- Software: B.D.P.
- Validation: B.D.P.
- Formal analysis: B.D.P.
- Investigation: B.D.P., M.W.
- Resources: M.W., J.T.V.
- Data curation: B.D.P., M.W.
- Writing - original draft: B.D.P.
- Writing - review and editing: B.D.P., M.W., C.E.P., J.T.V.
- Visualization: B.D.P.
- Supervision: C.E.P., J.T.V.
- Funding acquisition: B.D.P., C.E.P., J.T.V.

## 10 Acknowledgements

We thank Thomas Athey for helpful comments.

B.D.P. was supported by the [NSF Graduate Research Fellowship (Grant no. DGE1746891)](https://www.research.gov/grfp/AwardeeList.do?method=loadAwardeeList). J.T.V. was supported by the NSF CAREER Award (Grant no. 1942963). J.T.V. was supported by the NSF NeuroNex Award (Grant no. 2014862). J.T.V and C.E.P. were supported by the [NIH BRAIN Initiative (Grant no. 1RF1MH123233-01)](https://reporter.nih.gov/project-details/10012519).

## 11 Appendix

### 11.1 Understanding why BGM decreases accuracy for weak correlation

We sought to understand why bisected graph matching can decrease matching accuracy when the correlation in the contralateral subgraphs is weak. To do so, we repeated the simulation from Section 3 with a contralateral edge correlation of 0.1. Recall that for this value of the contralateral edge correlation, bisected graph matching achieves a matching accuracy of around 0.6, while for graph matching, accuracy is around 0.8. After simulating networks from this model, we computed the number of edge disagreements in the contralateral connections under four different permutations of the “right hemisphere”: the true alignment used to generate the network, the permutation predicted by graph matching, the permutation predicted by bisected graph matching, and the permutation predicted by “contralateral graph matching” (CGM, i.e., setting the ipsilateral networks to the zero matrix and running BGM).

For the parameters of the model used here, we observed that the true alignment of these networks had, on average, 26 edge disagreements (Figure 1). Note that for unweighted networks, the graph matching objective functions are trying to minimize the number of edge disagreements between the respective subgraphs being matched. The permutation predicted by graph matching yielded 27 edge disagreements – it is unsurprising that this is close to the true number, as graph matching is often recovering the true alignment as shown in Figure 2. However, we observed that bisected graph matching and contralateral graph matching yielded 23 and 16 edge disagreements on average, respectively. What should we make of the fact that there are actually *fewer* edge disagreements than under the permutation used to generate these subnetworks under the correlated Erdős-Rényi model?

This phenomenon was studied for correlated Erdős-Rényi networks in Lyzinski et al. [63], where the authors showed that if the correlation between the two networks is sufficiently weak, then the number of permutations which align the networks better than the true permutation goes to infinity as the networks grow. In other words, we can expect the performance of any graph matching algorithm to be poor (with respect to the true permutation) if the network correlations are weak. Given this phenomenon, [64] discussed how to assess these so-called “phantom alignments,” proposing methods for assessing whether a matching is likely to have arisen by pure chance under some model of uncorrelated networks. While these kinds of methods could also be applied to the bisected graph matching setting, they still do not provide a method for inferring a better matching for these weakly correlated networks, so we did not explore them further here.

### 11.2 Effect of network size in simulated experiments

In an extension to the simulation described in Section 3 we verified that the phenomenon shown in Figure 2 would hold over a range of network sizes. We used the same correlated Erdős-Rényi model described in Section 3, but varied the number of nodes in the simulated network. Specifically, we swept over a range of sizes which corresponded with the network sizes we studied for the real connectome datasets in Section 4.

Supplemental Figure 2 shows matching accuracy for GM and BGM as a function of network size, for contralateral correlations of 0.2, 0.4, and 0.6. We observed that again, when the contralateral correlation is too small (0.2), BGM performs worse than GM. However, for even modest correlation (0.4, 0.6), BGM provides a large improvement over GM across the entire range of network sizes. In fact, the gap between BGM and GM grows as the networks get larger. For instance, for networks with 600 nodes per side and a contralateral correlation of 0.6, GM is only matching with an accuracy of around 0.2, while BGM is matching perfectly.

**Supplementary Figure 1:**
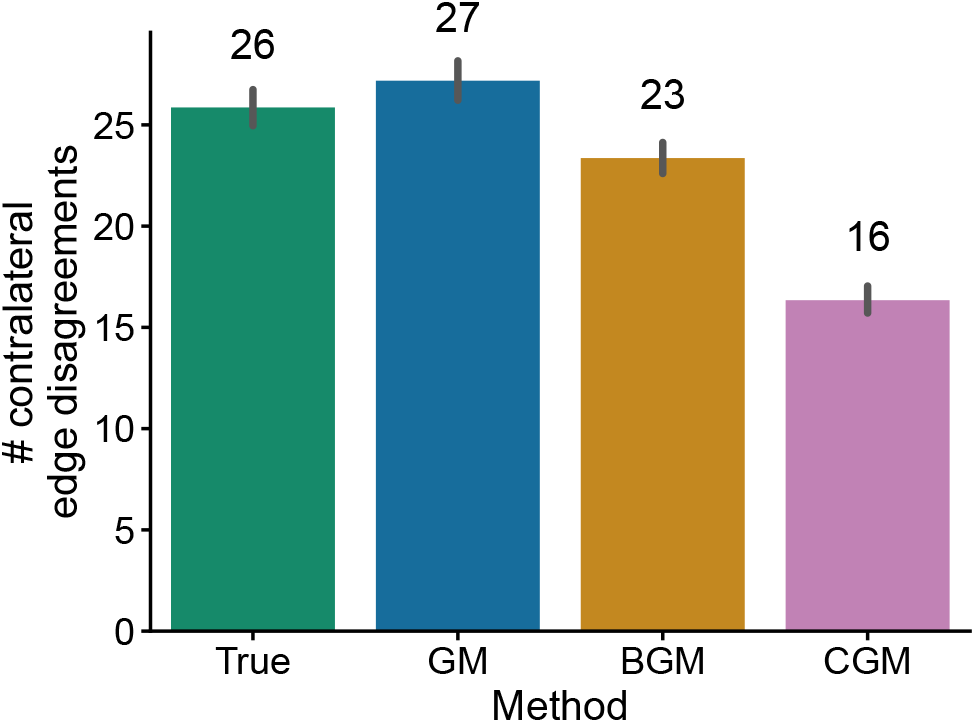
Examination of the number of edge disagreements after matching for the same network model described in Section 3 with a contralateral edge correlation of 0.1. Bars show the mean number of edge disagreements over 1,000 runs from the true alignment and the permutations predicted by graph matching (GM), bisected graph matching (BGM), and contralateral graph matching which ignores the ipsilateral subgraphs (CGM). Note that BGM and CGM are finding permutations under which there are fewer edge disagreements than under the alignment the contralateral networks were sampled from—in other words, they are finding permutations under which the contralateral networks are more similar to each other. This phenomenon explains why BGM has lower matching accuracy relative to the true permutation in Figure 2 when the contralateral correlation is very weak.

**Supplementary Figure 2:**
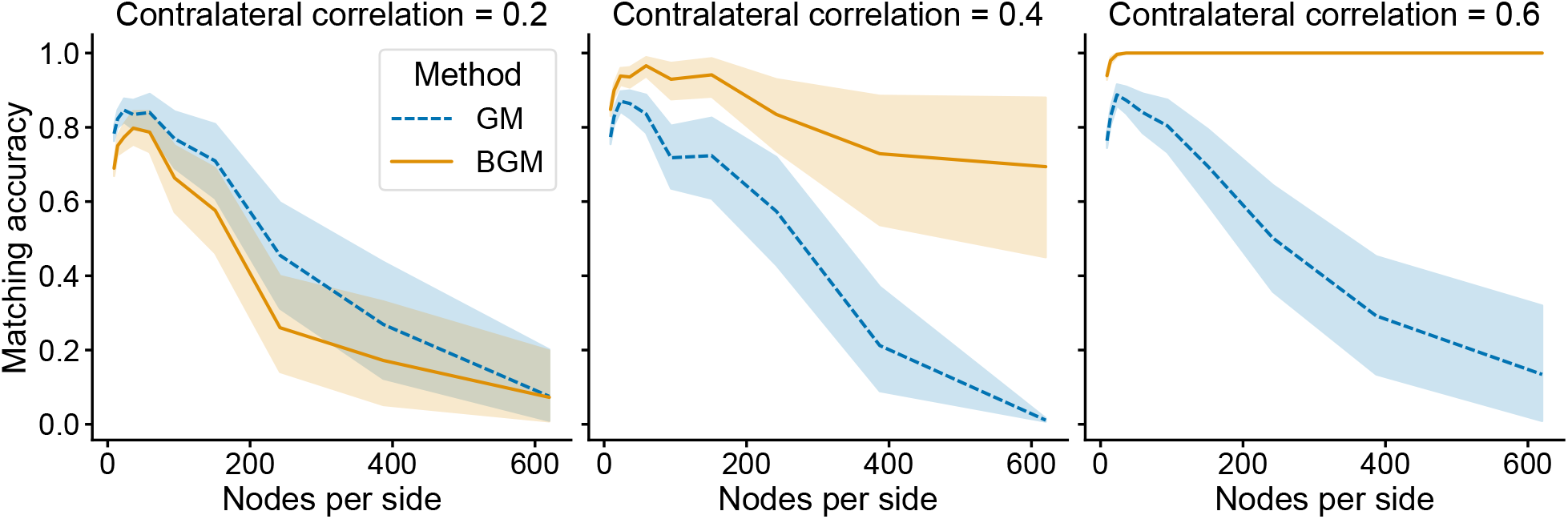
Comparison of matching accuracy for graph matching (GM) and bisected graph matching (BGM) as a function of network sizes, shown for varying contralateral correlations. Simulated networks were constructed following the same correlated Erdos-Renyi model as in Section 3, except the number of nodes (*n*) for each network was varied. As in Figure 2, when the contralateral edge correlation is too weak (0.2), then including the contralateral subgraphs in the optimization (BGM) does not improve performance for any network size. However, if the correlation is stronger (0.4 or 0.6 for this simulation), then BGM provides a large increase in matching accuracy across a range of network sizes.

### 11.3 Matching connectomes with simulated unpaired neurons

In many real connectome datasets, some subset of neurons may not have a homolog on the other side of the nervous system. Thus, we sought to understand the robustness of GM and BGM to the presence of these “unmatchable” nodes. To do so, we performed synthetic perturbations of the connectomes studied in Section 4. For each dataset, we randomly selected some proportion (ranging from zero to 0.25) of neuron pairs for perturbation. For these pairs, we shuffled the edges to and from these neurons, removing any edge correlations with their respective partners on the other side of the nervous system. Note that because some of these edges will be to/from the set of neurons which are still matched, this process will also make it harder to match those neurons as well. Then, we applied the graph matching and bisected graph matching algorithms to these perturbed connectomes, and measured the matching accuracy for the unperturbed neurons. This process was repeated 50 times for each dataset and percentage of neurons to perturb.

**Supplementary Figure 3:**
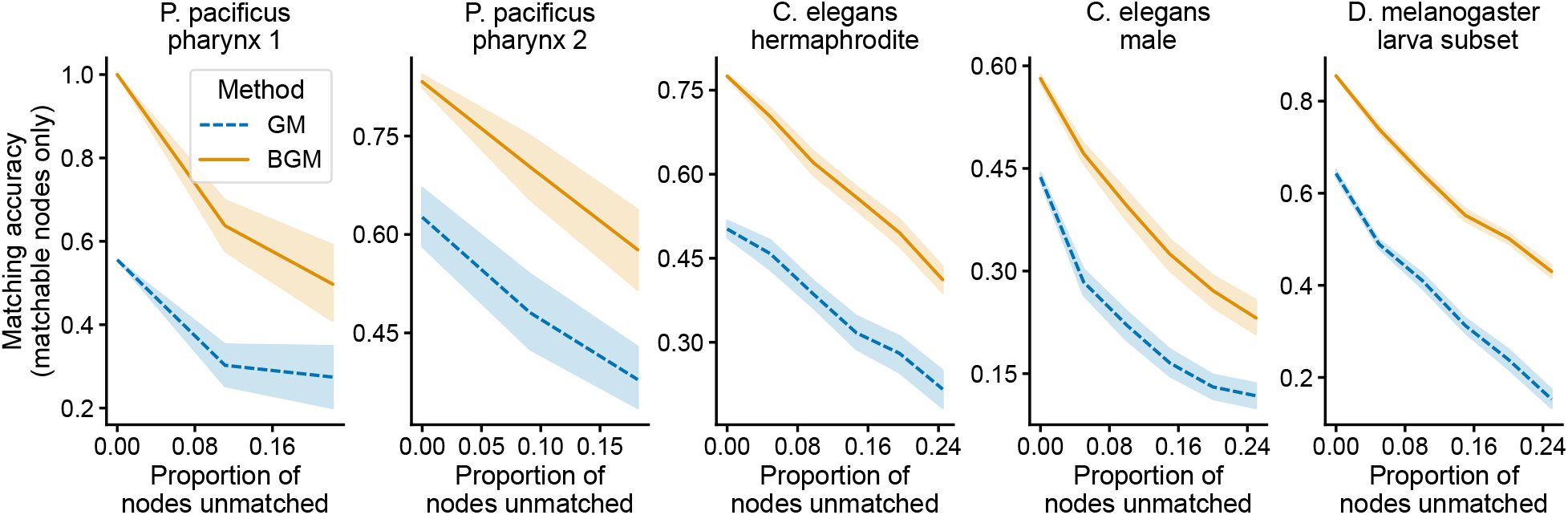
Mean matching accuracy across 50 simulations for graph matching and bisected graph matching for perturbations of the connectomes studied in Section 11.3. For each connectome (panels), some proportion (x-axis) of neuron pairs had their incident edges shuffled, removing any edge correspondence for those pairs. For each dataset and proportion of neuron pairs perturbed, bisected graph matching showed a higher mean matching accuracy (y-axis) for the remaining matchable nodes.

Figure 3 shows the matching accuracy for GM and BGM as a function of the proportion of pairs perturbed for all five connectome datasets. Unsurprisingly, unmatching some nodes by shuffling their incident edges does degrade the performance of both algorithms. However, we observed that in all cases, BGM had a higher matching accuracy than GM. Thus, BGM improves robustness in the presence of spurious nodes which have no true match on the other side of the nervous system.

### 11.4 Derivation of the adjusted gradient for bisected graph matching

Here, we show how the modified gradient (with respect to the permutation *P*) is computed for bisected graph matching. We start with the objective function for the bisected graph matching problem:

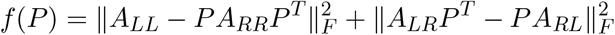

This can be written as the sum of a term for the ipsilateral subgraphs (*f*_*I*_(*P*)) and one for the contralateral subgraphs (*f*_*C*_(*P*)):

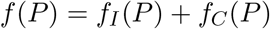

Rewriting the first term:

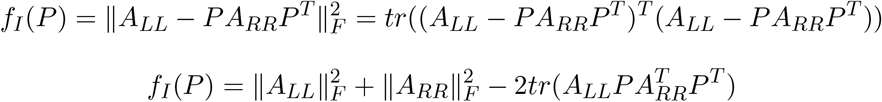

And likewise for the second:

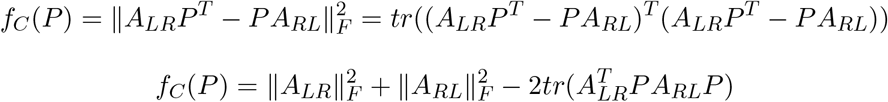

We can drop terms unrelated to *P* and constants (since they will not affect the optimization) and returning to the full objective function we have:

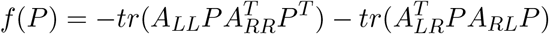

Then, taking the derivative with respect to *P* for the first term:

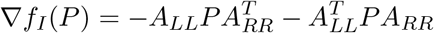

Similarly for the second term:

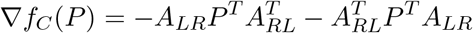

So for the full objective function, the gradient is:

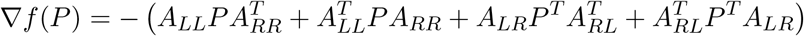

**Supplementary Figure 4:**
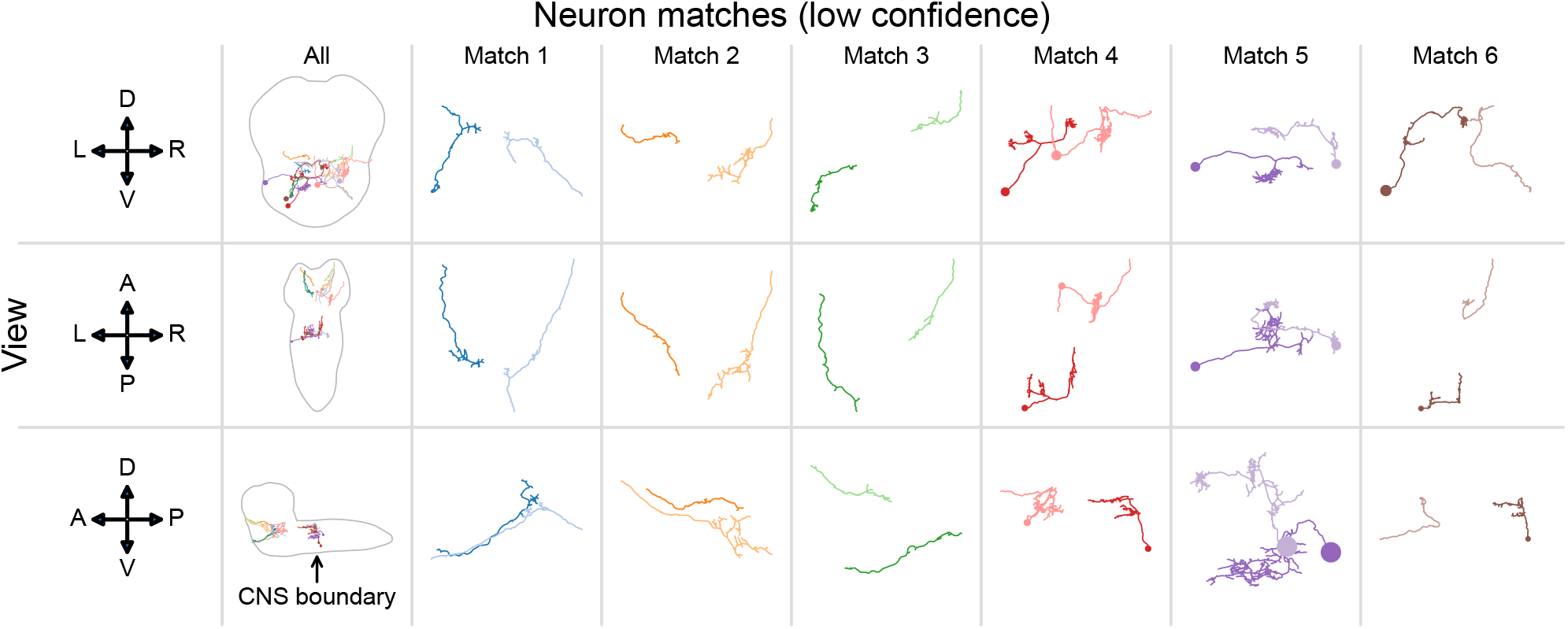
Morphological comparison of low-confidence matched neurons in the *Drosophila* larva connectome subset [8–28] (see Section 5.2). Each column shows the best match for a left-hemisphere neuron which was frequently matched to different neurons across BGM initializations. Thus, these neurons often change partners over each initialization of the algorithm, suggesting that they are unlikely to be true bilateral homologs. Each row shows a different view of a matched pair of neurons (anatomical axes to the left show: D-dorsal, V-ventral, L-left, R-right, A-anterior, P-posterior). The morphology of these matched neurons appears different in most cases (compare to Figure 6), further supporting the idea that these matches represent implausible candidates for true bilateral homologs.

